# Virus specific impacts on honey bee flight performance are mediated by the octopamine pathway

**DOI:** 10.64898/2026.03.11.710992

**Authors:** Naomi G. Kaku, Michelle L. Flenniken

## Abstract

High annual honey bee colony losses are associated with environmental and biological stressors, including virus infections. In insects, the octopamine pathway orchestrates the “fight-or-flight” response, regulating energy mobilization, temperature, and flight. We determined that sacbrood virus (SBV) infections induce expression of an octopamine receptor and enhance honey flight performance, whereas deformed wing virus (DWV) infections reduce flight performance, but how viruses interface with this pathway remained unknown. To elucidate the relationships between the octopamine response, virus infection, and flight, honey bees were infected with SBV or DWV and exposed to octopamine (OA), epinastine (EP)–an OA receptor antagonist, or both OA and EP; flight and gene expression were assessed. Pharmacologic manipulation revealed that octopamine supplementation rescued flight deficits in DWV-infected bees, but diminished performance in SBV-infected bees, while blocking octopamine receptors altered these effects. Transcriptome analyses indicated that SBV infections, and DWV infection with OA treatment, activated honey bee metabolic pathways, and that SBV infected bees had greater expression of genes involved in OA synthesis, unless treated with OA. These results provide mechanistic insight for virus-specific impacts on honey bee flight, which may have consequences on foraging efficiency, colony health and virus transmission.

**Impact Statement:** Differential virus-specific impacts on honey bee flight performance involve octopamine signaling and chemical stressors affecting this pathway may impact colony health.

## Introduction

Honey bees (*Apis mellifera*) are eusocial, generalist pollinators that live in densely populated colonies of ∼30,000 individuals (1–3). The survival of each colony depends on the coordinated activities of thousands of individual bees that provision nutritional resources, rear brood, and regulate internal hive temperature (3–6). Colony losses in the U.S. averaged 40% from 2008-2025 (7–11), these losses are associated with several abiotic and biotic stressors, including parasites and pathogens (12–16). Honey bee pathogens include bacteria, microsporidia, and viruses (17, 18). Viruses may be transmitted vertically, horizontally, or by vectors, including the *Varroa destructor* mite (19–23). *Varroa* mites feed on honey bees and infestations may kill colonies (24). In addition, *Varroa* mites vector viruses, including deformed wing virus (DWV, *Iflavirus aladeformis*), which replicates in the mite, and others that are mechanically transmitted such as sacbrood virus (SBV, *Iflavirus sacbroodi*) (25–27). Most characterized bee infecting viruses have positive single-stranded RNA (+ssRNA) genomes including members of *Iflaviridae* (e.g., DWV and SBV). Virus infections in honey bees may remain seemingly asymptomatic, cause morphological symptoms, paralysis, and/or result in death (17, 28–32). Honey bees that lack morphological symptoms may still harbor virus infections that impact lifespan and flight performance (8, 33–39).

Flight is an energetically taxing behavior that is essential for honey bee foraging, defecation, and mating (40–45), and flight muscles are essential for temperature regulation (i.e., fanning to cool and ‘shivering’ for heat) (19, 46–49). In our previous work, we used flight as a proxy for overall bee health and demonstrated that honey bees harboring high levels of DWV without obvious symptoms had impaired flight performance (33). Unexpectedly, we determined that the flight performance of SBV infected honey bees was enhanced and linked to the metabolism-stimulating, stress-induced octopamine pathway (33). Abiotic and biotic stressors may induce enhanced activity of stress pathways including the octopamine response. Invertebrates synthesize octopamine (OA), a ‘fight or flight’ neurohormone analogous to norepinephrine, which facilitates energy production for taxing activities including flight (50–54). The octopamine response may also be triggered by other stressors, including pathogen infections (33, 55, 56).

Whether octopamine signaling is immunosuppressive or immunoenhancing is dependent on multiple factors and its role in honey bee antiviral defense has not been thoroughly investigated (57, 58). Honey bee behaviors may be influenced by multiple monoamine neurohormones including OA, serotonin, dopamine, and tyramine (51, 59–64). Octopamine promotes movement, neuromuscular signaling, and metabolism. To synthesize OA, tyrosine is first converted to tyramine by Tyrosine decarboxylase (TDC); tyramine is the precursor neurohormone to OA and is generally associated with reduced flight and movement (52, 65, 66). Tyramine may serve as an antagonistic modulator of behavior to OA or Tyramine β-hydroxylase (TβH) may convert it to OA (Fig 1A). One of the most abundant OA receptors in honey bees is a G-coupled protein receptor, Oβ-2R (67). When OA binds to Oβ-2R, it activates adenylyl cyclase (AC), which increases intracellular concentration of cyclic adenylyl monophosphate (cAMP). Elevated cAMP activates cAMP-dependent Protein kinase A (PKA) which phosphorylates metabolic enzymes and transcription factors that generally enhance metabolic activity (Fig 1B) (47, 53, 67, 68).

**Figure 1.**
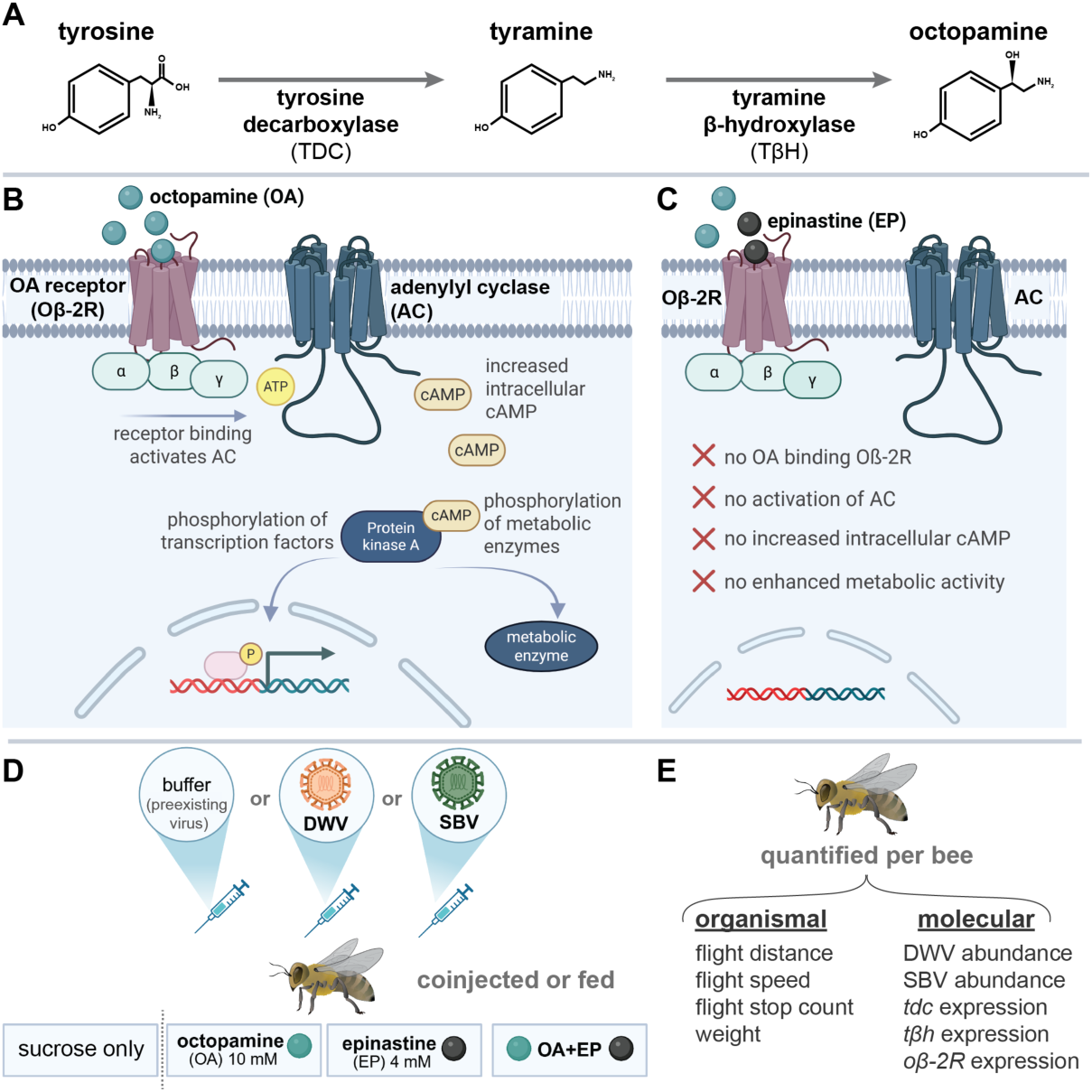
Octopamine signaling pathway (i.e., synthesis, activation, inhibition) and experimental design. (A) Octopamine (OA) is synthesized by conversion of tyrosine to tyramine by tyrosine decarboxylase (TDC) and tyramine conversion to OA by tyramine β-hydroxylase (TβH). (B) OA β-2 receptor (Oβ-2R) is one of the most abundant and specific OA-detecting receptors. Upon activation of Oβ-2R, a G-coupled protein receptor, adenylyl cyclase (AC) is activated and increases intracellular cyclic adenylyl monophosphate (cAMP) levels. High cAMP levels induce protein kinase A activity which phosphorylates metabolic enzymes and transcription factors that enhance metabolic activity. (C) An OA receptor antagonist, epinastine (EP), inhibits OA binding and prevents activation of the Oβ-2R pathway. (D) To evaluate relationships between virus infection and the OA response, honey bees were inoculated with either deformed wing virus (DWV) or sacbrood virus (SBV) via injection or mock infected. Bees were exposed to either OA, EP, or both OA and EP via co-injection or feeding, all bees were fed sucrose. (E) Impacts of experimental conditions were quantified in individual bees 72 hours post infection. Panels B-D produced in BioRender (Kaku, 2026).

We hypothesized that SBV infections, which correlate with higher expression of an OA receptor (Oβ-2R) (33), activate the OA response which results in enhanced flight performance. Therefore, we predicted that OA treatment would enhance honey bee flight performance regardless of virus infection. Epinastine (EP) is a highly specific OA receptor antagonist, exhibiting at least four orders of magnitude greater affinity for OA receptors than for other honey bee biogenic amine receptors (69, 70). Therefore, we hypothesized that EP treatment would reduce honey bee flight performance and negate SBV-associated enhanced flight performance.

## Results

To test these hypotheses, we exposed virus infected honey bees to OA, EP, or both OA and EP and assessed flight performance and gene expression (e.g., *tdc*, *tβh*, and *oβ-2R*) in three independent experiments (i.e., exp-1, 2, and 3) (Fig 1 and Data S1). Specifically, age-matched, one day old adult bees (n = 336) were experimentally infected with DWV (n = 117), SBV (n = 75), or mock infected (buffer injection, n = 144) and treated with OA (10 mM), EP (4 mM), or both OA and EP via injection or feeding, or not treated; all bees were fed 50% sucrose syrup *ad libitum*. Treatments of OA and EP were given in 2 µL injections or fed at the same concentration in sucrose syrup; doses were in-line with previous studies (33, 47, 52, 69). Exposure to OA and/or EP via injection was temporally limited, whereas *ad libitum* feeding provided continuous exposure (Fig 1).

### Quantification of experimentally introduced and naturally occurring honey bee virus infections

Honey bees used in this study were sourced from managed colonies that are naturally exposed to pathogens. To limit variation associated with confounding infections, bees were collected upon emergence under laboratory conditions (19, 33, 71, 72). Therefore, in addition to the use of quantitative polymerase chain reactions (qPCR) to determine the abundance of DWV and SBV in individual honey bee samples, PCR was used to test for potential preexisting infections and detected pathogens were quantified using qPCR, as described previously (33) (Data S1-S3). We determined that honey bees in exp-1 had preexisting DWV infections and bees in exp-2 and exp-3 had preexisting infections of DWV and SBV (S1 Fig). Most bees analyzed herein had either experimentally introduced or preexisting DWV infections (n = 325/336) and 40% had experimentally introduced or preexisting SBV infections (n = 134/336). DWV infections ranged between 1×10^3^ and 7.1×10^10^ copies per 2 µg of RNA (∼7×10^3^ – 8×10^11^ DWV copies per bee). SBV infections were between 1.2×10^3^ and 1.2×10^10^ copies per 2 µg of RNA (∼8.3×10^4^ – 8.3×10^11^ SBV copies per bee). Due to high DWV and SBV prevalence there were no virus-free honey bees. Mock infected bees harbored low virus levels (i.e., an average of 3×10^5^ DWV copies and 2×10^3^ SBV copies per 2 µg RNA) and inoculation resulted in higher virus loads (i.e., an average of 2×10^8^ DWV copies and 2×10^8^ SBV copies per 2 µg RNA), *p* = 2×10^-16^ and 5×10^-11^, respectively). To compare the impact of varying virus levels and treatments, we evaluated several flight metrics (i.e., flight distance, duration, speed, and stop count; Data S1) using linear mixed effect models (LMMs) (complete information for each model and model selections in methods and Data S4).

### OA and EP altered flight performance of virus infected bees

The previously described DWV-associated flight impairment relative to mock infected bees (33), was recapitulated in this study (i.e., DWV infected bees flew shorter distances (*p* = 0.005)). Mock infected bees from this sample cohort, which harbored low virus levels, flew an estimated 45 meters, whereas bees harboring high DWV levels (i.e., > 10^8^ DWV RNA copies / 2 µg RNA) flew only ∼9 meters. In line with our hypothesis, DWV-associated flight impairment was alleviated by OA feeding or injection (i.e., average flight distance was similar to mock infected bees fed sucrose, *p* = 0.61 and 0.08, respectively) (Fig 2A). Honey bees that received EP flew shorter distances (*p* = 0.002), but the effect of feeding EP was not assessable since bees with experimentally introduced DWV infections that were fed EP died, and there were only 14 bees with low DWV infection levels (Fig 2B, Data S4). Co-injection of OA and EP negatively impacted the distance DWV infected honey bees flew (*p* = 0.020) (Fig 2C). Counter to our hypothesis, OA-treatment did not enhance flight performance of SBV infected bees (*p* = 0.69 and 0.60) (Fig 2D, Data S4). SBV infected bees that were treated with EP or co-treated with both EP and OA via injection flew shorter distances than untreated SBV infected bees (*p* = 0.022 and 0.004, respectively), whereas feeding had no impact (Fig 2E-F).

**Figure 2.**
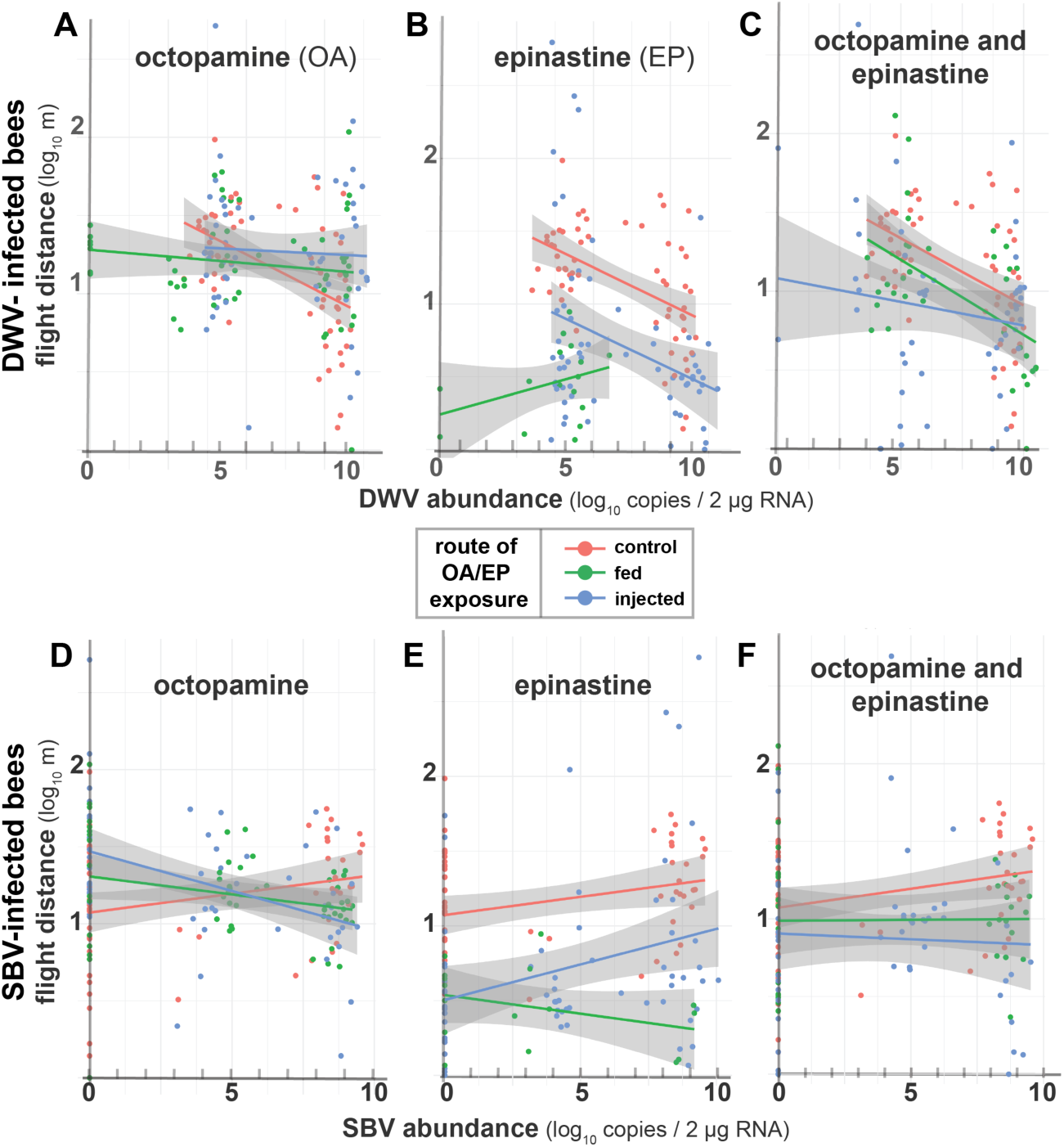
The flight performance of honey bees infected with DWV and/or SBV is differentially impacted by OA and EP treatment. To evaluate the involvement of the OA pathway on virus infection and/or flight performance, virus-or mock-infected honey bees were exposed to OA, EP, or both OA and EP via co-injection (blue) or feeding (green). Honey bees were experimentally infected with or had preexisting infections of DWV (A-C) or SBV (D-F) in three experiments. Flight distance (log_10_ m) was evaluated using linear mixed effect models to quantify impacts of DWV and/or SBV infection in the context of OA and/or EP treatment. Each data point represents an individual bee (n = 336); colors indicate treatment (i.e., bees fed only sucrose in pink, bees fed OA and/or EP in green, and bees injected with OA and/or EP in blue) and gray areas represent 95% confidence intervals. Linear mixed models with estimates in Data S4. (A) In bees fed only sucrose, DWV infections negatively impacted flight distance (*p* = 0.005), whereas DWV infected bees treated with OA flew further distances (i.e., average flight distance was similar to mock infected bees fed sucrose, *p* = 0.61 and 0.08, respectively). (B) DWV infected bees injected with EP flew shorter distances than DWV infected bees fed only sucrose (*p* = 0.002). (C) Co-injection of OA and EP negatively impacted the distance DWV infected honey bees flew (*p* = 0.020) (D) OA-treatment did not enhance flight performance of SBV infected bees (*p* = 0.69 and 0.60) (E) Honey bees with SBV infections fed or injected with EP flew shorter distances than SBV infected bees fed only sucrose (*p* = 0.022 and 0.004, respectively). (F) Bees with SBV infections fed or injected with OA and EP resulted in similar flight distances regardless of SBV infection level.

The amount of time a honey bee was actively flying was also assessed as an indicator of energetic capacity (S2 Fig). Honey bees harboring high DWV levels (i.e., >10^8^ DWV copies / 2 µg RNA) flew only 14 minutes (*p* = 0.022), relative to mock infected bees with low virus infection levels (average of 3×10^5^ DWV copies / 2 µg RNA), which flew for 26 minutes. Honey bees with high SBV levels flew for similar durations to mock infected bees (i.e., 28 minutes, *p* = 0.58), which is consistent with previous results (33). In line with our hypotheses, mock infected bees with low virus infection levels flew longer when they were fed OA (i.e., 49 minutes) (*p* = 0.0001). DWV-associated flight impairment was mitigated in honey bees that were supplemented with OA, which flew for similar durations to mock infected bees (i.e., 28 minutes, *p* = 0.76). However, counter to our hypothesis, when SBV infected bees were injected or fed OA, flight duration was significantly reduced (i.e., 18 and 12 minutes, respectively; *p* = 0.04 and 0.0009).

To quantify the impact of virus and OA and/or EP treatment on honey bee flight speed, we compared average and peak speeds. Average flight speed (i.e., total distance / total flight duration) was similar across all conditions except in bees fed EP, which flew at slower average speeds (*p* = 3×10^-4^) (Data S4). We also compared peak speed (i.e., maximum speed reached during flight sustained >10 seconds) (S2 Fig). Similar to previously reported results (33), DWV infected bees had slower peak speeds than mock infected bees (*p* = 0.016). SBV infected bees flew similar speeds to mock infected bees (*p* = 0.07), but peak speeds were reduced when SBV infected bees were fed or injected with OA (*p* = 0.004, *p* = 0.0006) (Data S4). Mock infected bees flew at peak speeds of ∼3.3 km/h, honey bees with high levels of SBV (i.e., 10^8^ SBV copies / 2 µg RNA) flew ∼3.8 km/h, whereas highly SBV infected bees injected or fed OA flew an estimated 2.0 or 2.5 km/h, respectively (*p* = 0.001 and 0.022).

In addition to flight metrics, post-flight thorax temperatures were measured within a subset of individual bees (n = 130) since flight induces heat and the octopamine response is involved in honey bee temperature regulation (47, 67) (Data S1 and S4). The ambient temperature for all flight assays was 22°C, and post-flight thorax temperatures in honey bees maintained in control conditions averaged 28°C. DWV infected bees had similar post-flight temperatures, and highly SBV infected bees were 2°C cooler than control (*p* = 0.04). Honey bees with DWV infections that were fed OA had thorax temperatures that were 2.5°C higher than bees not fed OA. Whereas the thorax temperatures of bees with DWV infections were 5.6°C lower in bees co-fed OA and EP (*p* = 0.027 and 1×10^-6^). Likewise, honey bees with high SBV levels that were injected with OA also had reduced thorax temperatures post-flight (i.e., ∼ 3°C lower) (*p* = 0.03; Data S4).

### Virus specific outcomes of experimentally manipulated octopamine pathway explained by differential expression of key genes

Induction of the OA pathway was previously implicated in the enhanced flight performance of SBV infected honey bees, which had elevated expression of *oβ-2R* (33). When OA binds to Oβ-2R, it activates AC, increases cAMP concentration, thereby activating cAMP-dependent PKA, which phosphorylates enzymes and transcription factors that enhance metabolic activity (Fig 1B) (47, 53, 67, 68). To determine if DWV, SBV, or different experimental treatments were associated with differences in OA synthesis, we quantified the expression of *tdc* and *tβh* relative to mock infected honey bees fed only sucrose (Data S1, Fig 3). We evaluated the relationship between *tdc* expression and multiple factors including DWV abundance, SBV abundance, experimental treatments (i.e., OA and/or EP, fed/injection) using LMMs which explained 91% of the data variation (Data S4). SBV infection was associated with increased *tdc* expression (*p* = 0.021), and SBV infected bees fed OA or co-fed OA and EP also expressed higher *tdc* levels (*p* = 0.030 and 2×10^-10^, respectively). Expression of *tdc* was lower in SBV infected bees co-injected with EP, and SBV and DWV-coinfected bees that were fed OA or both OA and EP (*p*-values = 0.033, 0.022, 0.006, 0.004, respectively) (Fig 3A, Data S4). Expression of *tdc* was not associated with DWV infection, or DWV infection with OA and/or EP treatment (*p*-values all >0.2, Data S4, S3 Fig). We also examined the effects of virus infection in the context of experimental OA and EP treatments on *tβh* expression (i.e., in bees with detectable *tβh* levels n=306/336) (Fig 3B). An LMM that included DWV abundance, SBV abundance, treatment, and *tdc* expression as explanatory variables explained 85% of the variation in *tβh* expression (Data S4). Expression of *tdc* and *tβh* were positively correlated (with a *tdc*:*tβh* ratio of 1:1.6, *p* = 0.004) (S4 Fig). Expression of *tβh* was higher in SBV and DWV infected bees (*p* =0.022 and 1×10^-9^, respectively), suggesting more tyramine is converted to OA during DWV or SBV infection (Fig 3B). Results from these models indicate that only SBV infections were associated with greater expression of both *tdc* and *tβh* relative to mock infected bees with low preexisting virus levels. Whereas SBV infected honey bees that were co-injected with OA expressed similar levels of *tdc* or *tβh* as mock infected bees (*p* = 0.421 and 0.437, respectively). This result suggests that the elevated expression of genes involved in OA synthesis in SBV infected honey bees is reduced by OA supplementation and partially explains the impaired flight performance of these bees relative to untreated SBV infected honey bees (Fig 4). For OA to stimulate metabolic activity and energy production, OA must activate receptors including Oβ-2R (67). To identify variables that contribute to the range of *oβ-2R* expression, we compared LMMs including *tβh* expression, *tdc* expression, SBV abundance, and OA/EP treatment, which explained 95% of the data variation (S4 Fig). We considered including DWV abundance as a fixed effect in this model, but it did not appreciably contribute to *oβ-2R* expression (Data S4). These results corroborate our previous findings that SBV, but not DWV, was associated with increased *oβ-2R* expression (33). The strongest predictors of *oβ-2R* expression levels were *tdc* and *tβh* expression (*p* = 4.4×10^-5^ and 3.9 x 10^-5^) and SBV was positively associated with *tdc* and *tβh* expression (*p* = 0.0006) (Data S4).

**Figure 3.**
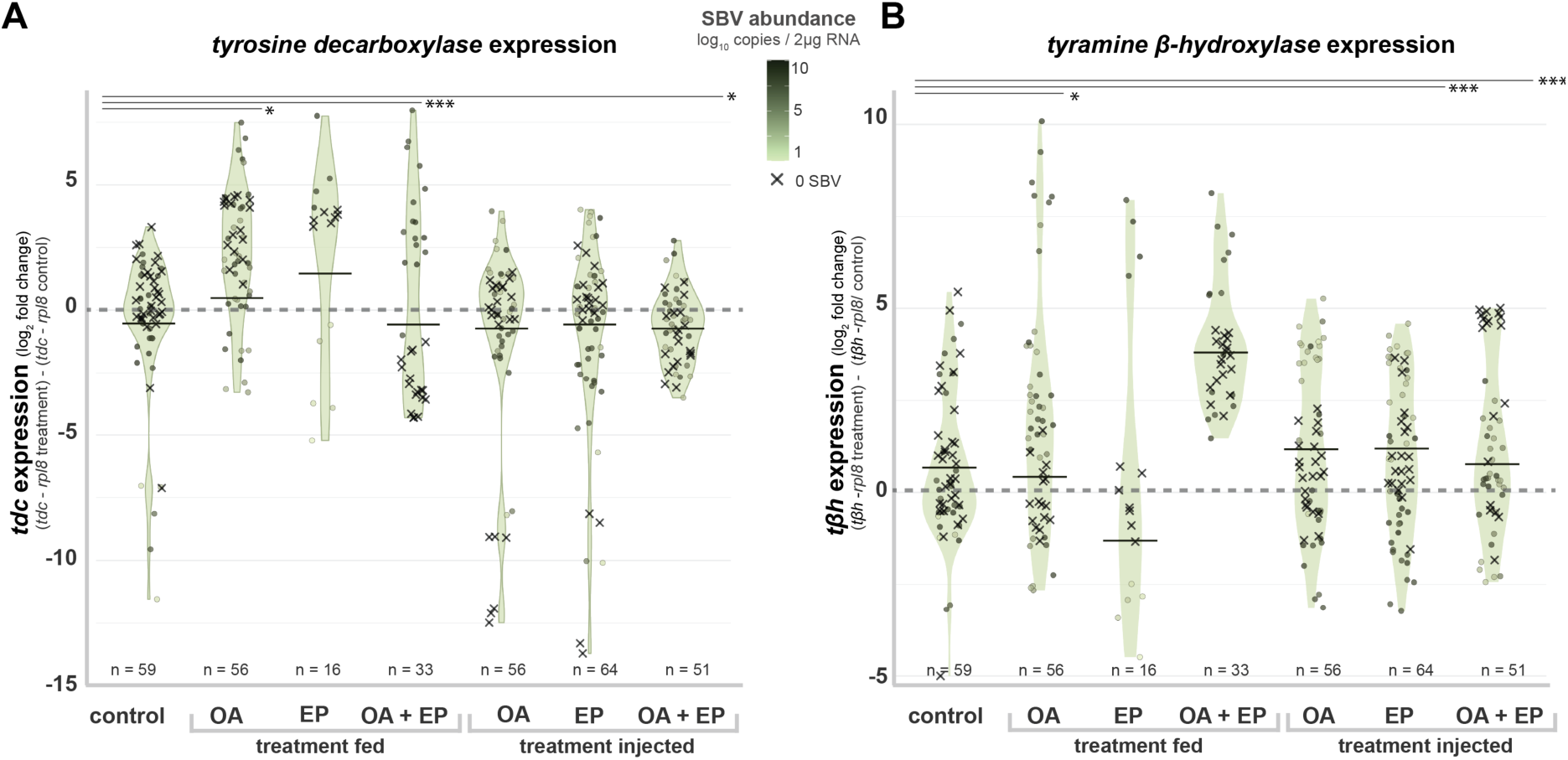
Expression of *tyrosine decarboxylase* (*tdc*) and *tyramine β-hydroxylase* (*tβh*) across treatments and SBV infection. The relationships between honey bee *tyrosine decarboxylase* (*tdc*) and *tyramine β-hydroxylase* (*tβh*) expression in the context of virus infection and treatments were evaluated using a linear mixed effect model that included SBV abundance and OA/EP treatments as fixed effects with an interaction. Individual honey bee data plotted by treatment group with either (A) *tdc* expression or (B) *tβh* expression. Gene expression was calculated as log_2_ fold change using the ΔΔCt method relative to housekeeping *rpl8* and compared to mock infected bees. SBV abundance was calculated as log_10_ copies / 2 µg RNA. The background violin plots represent the 95% confidence interval, and the horizontal line in each violin represents the median. SBV abundance is represented by a green color scale and SBV-negative samples with an “X”. When SBV abundance did not impact *tdc* or *tβh* expression the dark green points are evenly distributed above and below the control ‘0’ fold change line, whereas unequal distribution indicates an SBV-specific effect; * p<0.05, **p<0.01, and ***p<0.001. (A) Positive association between SBV abundance and *tdc* expression (*p* = 0.021). SBV infected bees fed OA or both OA and EP had greater *tdc* expression levels than uninfected bees fed only sucrose (*p* = 0.03 and 2×10^-10^). (B) Expression of *tβh* was greater in SBV infected bees than uninfected bees (*p* =0.022). SBV infected bees fed OA had greater *tβh* expression levels than uninfected bees fed sucrose (*p* = 2×10^-5^). SBV infected bees injected with EP or co-injected with OA and EP had lower *tβh* expression (*p* = 2×10^-5^ and 4×10^-9^).

**Figure 4.**
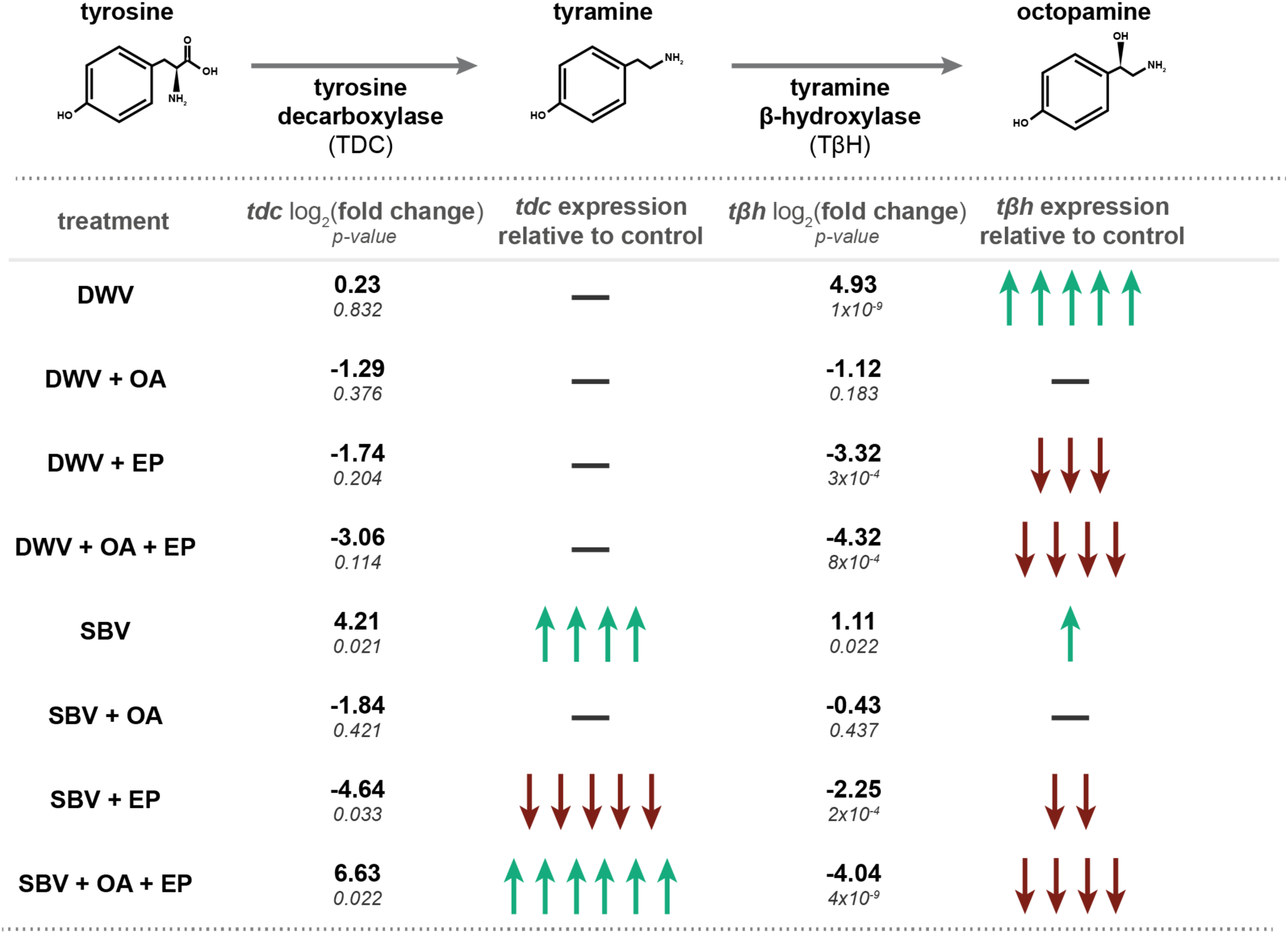
Relative *tdc* and *tβh* expression in honey bees with high DWV or SBV levels. (A) Tyrosine is converted via tyrosine decarboxylase (TDC) to tyramine. Tyramine is a neurohormone that acts as a behavioral antagonist to octopamine (OA), resulting in reduced movement in invertebrates. Tyramine may act as a signaling neurohormone, or tyramine β-hydroxylase (TβH) may convert it to OA. (B) A linear mixed model was used to assess the relative expression of *tdc* and *tβh* in honey bees harboring high levels of either deformed wing virus (DWV) or sacbrood virus (SBV) (i.e., 108 virus copies / 2μg RNA) in the context of OA and/or epinastine (EP) injection. Estimates of relative *tdc* and *tβh* expression were generated based on data from 336 honey bees. The single bold dash indicates *tdc* or *tβh* expression was similar in virus infected bees and mock infected bees and arrows indicate estimates that were higher or lower than controls where each arrow represents fold change of 1.

### Transcriptome level impacts of virus infection, flight, and OA treatment

We hypothesized that flight-impacting virus infections would result in differential expression of genes involved in stress, metabolic, and immune pathways. To test this, the transcriptomes of SBV, DWV, DWV and OA, or mock infected honey bees were compared. Honey bees were obtained from a previous experiment and included individuals that did or did not fly (33). RNA was prepared for high throughput sequencing on an Illumina NovaSeq X Plus 25B (n = 36 total individual bee samples, 3-5 per treatment group; Data S6). An average of 25 million reads was obtained for each sequencing library and mapped to the *A. mellifera* assembly HAv3.1 (73, 74). To evaluate similarities and differences between the transcriptional responses of honey bees with flight-impacting virus infections, we compared DWV, SBV, and DWV and OA injected bees relative to mock infected bees immediately post-flight since these bees had little to no detectable preexisting infections, whereas the no-flight mock infected bees had preexisting DWV infections (Data S5-S6).

Hundreds of differentially expressed genes (DEGs) were identified in SBV, DWV, and OA treated DWV infected bees (Fig 5). DWV infected bees with and without OA treatment shared 95 DEGs, which was more than they shared with SBV infected bees, indicating that there are virus specific transcriptional responses. There were 24 genes shared among SBV, DWV, and DWV and OA injected bees including elevated expression of those involved in the Ras-Raf-MEK-ERK pathway (including *mesr3* and *rap2l*) which is associated with metabolism, muscle development, and the cAMP/PKA pathway (75). Genes with lower expression were involved in muscle function (i.e., *mlc2, mlc1, cher, sar*), energy production (i.e., *cnn, ant, pka-c3*), and an inhibitor of proteolytic activity (i.e., *cyp4g11*).

**Figure 5.**
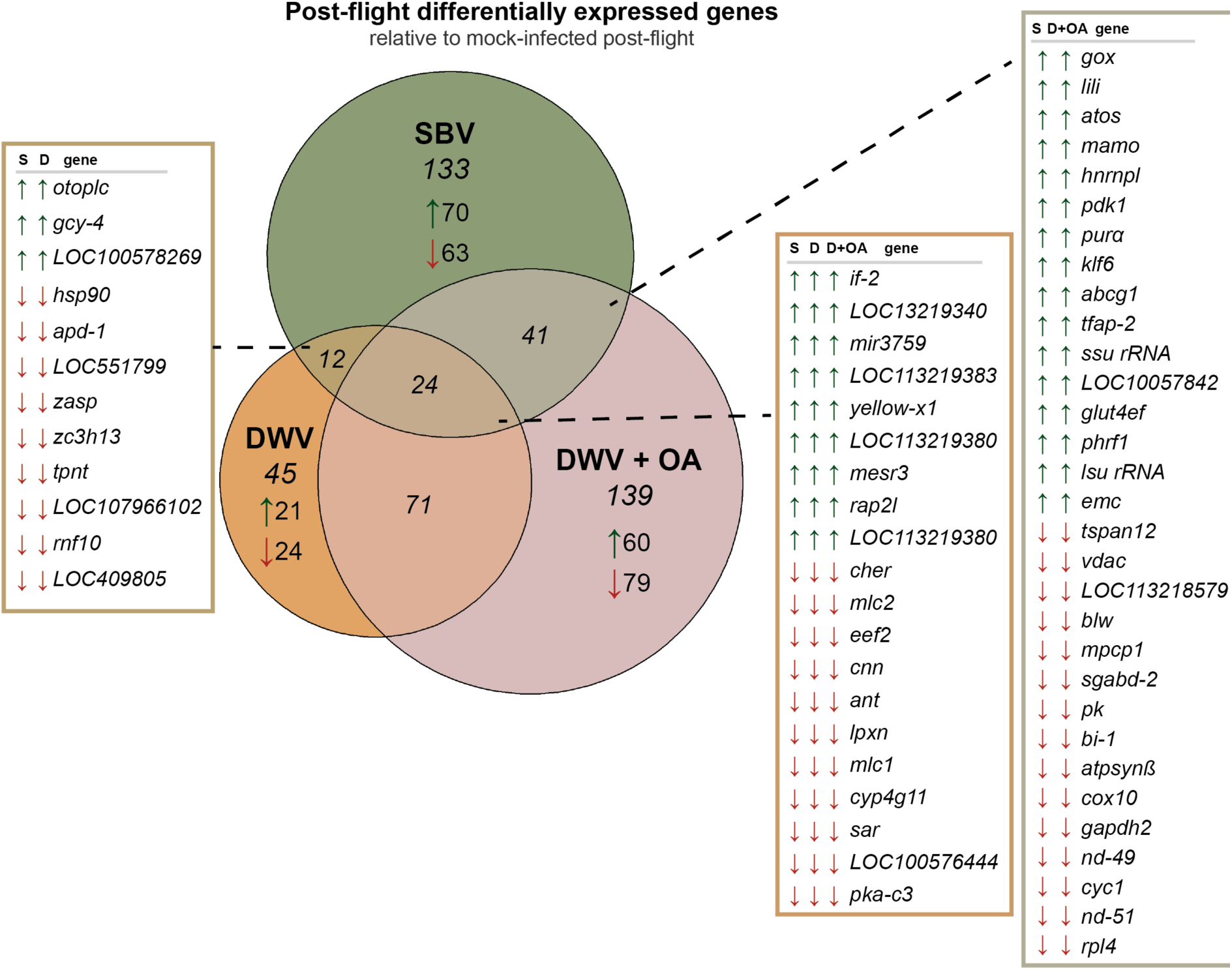
Transcriptome level comparison of honey bees that were SBV infected, DWV infected, or DWV infected and OA-treated. Gene expression was evaluated in honey bees that were mock infected, SBV infected, DWV infected, or DWV infected bees that were co-injected with OA (DWV+OA) after flight. Differentially expressed genes (DEGs) for each treatment group were identified relative to mock infected bees and compared using a Venn diagram. Full list of DEGs, fold change, and Benjamini-Hochberg corrected significance values are included in Data S6.

There were many genes unique to SBV and DWV infections (Fig 5). Honey bees with SBV infections exhibited lower expression of several genes associated with the citric acid cycle (*acsl6*, *cg5065*, *acyl-coA delta*(*11*), *mcad*, *nd-13a*, *nd-13b*, *nc73ef*, and *mdh2*) and general muscle function and tissue repair genes (*tmp-1, tmod, plastin-2, and fgf17*). SBV infected bees had elevated expression of *rgs11*–which increases G-protein α-subunit active state to inactive to expedite the G-coupled protein response (76), *titin*–which contracts stretched/striated muscle tissues (77), and several immune genes such as *argonaute-2*, *relish*, and *domeless* (Data S6-S7). DWV infected bees exhibited greater expression of several genes associated with immune function including *protein mahjong*, *spastin*, and a small heat shock protein, *protein-lethal*(*2*) *essential for life-like* (Data S6). The DEGs unique to DWV infected bees post-flight that exhibited lower expression included genes involved with muscle function and structure (*obscurin*, a titin homolog, *ryr*, *actin*, *mhc*, *stacl*, *tpm1*, and *zasp66*) and ATP generation (*citrate synthase, adenylate kinase 1, ADP/ATP translocase*).

Since OA supplementation mitigated DWV-associated flight impairment, we identified differences associated with OA by comparing DEGs to untreated DWV infected honey bees post-flight. As hypothesized, DWV infected bees that were co-injected with OA treatment had greater expression of OA pathway genes (*oβ-2R*, *adenylyl cyclase (ac)*, and CREB binding proteins) and genes involved in glucose metabolism (*gale*, *gdh*, *adpgk*, *putative glucosylceramidase 4*, *gba1lp, g6pd*, *sugarless*, *pdp*, *ND-24*, *NAD kinase*, *tret1*, *1,5-anhydro-D-fructose reductase, and igfals*) (Data S8-S9). These bees also expressed higher levels of genes induced by high cyclic guanine monophosphate (cGMP) levels (i.e., *burgundy*, *cGMP-dependent protein kinase-like*, and *cGMP-dependent protein kinase 1*) and genes in the RAS/MAPK-pathway (*rho1*, *rac1*, and *rab5*). Untreated, DWV infected bees expressed higher levels of several immune genes (*abaecin, domeless*, *echinoid*, and *defensin-1*) and heat shock protein encoding genes (*hsp83*, *hsp90*, *hspβ-1*) relative to DWV infected bees treated with OA (Data S7-S9).

After flight, both SBV infected and OA treated DWV infected bees expressed lower levels of several mitochondrial associated genes (*vdac, blw, atpsynβ*, *cox10*, *klf6,* and *mcmp1*) and genes involved in energy generation (*pyruvate kinase*, *gapdh2,* and 2 *ND-49,* and *ND-51*) than mock infected bees (Fig 5). This result suggested that flight was more taxing for virus infected bees. In addition, transcriptome level results indicated that OA treatment during DWV inoculation resulted in enhanced energy production, since these bees expressed higher levels of genes involved in energy production post-flight relative to DWV infected bees that did not fly (Data S10).

Since OA treatment resulted in recovered flight performance in DWV infected bees, we compared the transcriptomes of honey bees with similar DWV-levels that were either OA-treated or untreated. The 50 genes with the greatest difference in mean expression were identified in bees that did or did not fly (i.e., 100 total possible genes) (Data S10). There were 30 shared genes associated with OA-treatment regardless of flight status. This list included *defensin-1*, *apidermin-2*, *melittin*, *odorant binding protein 3*, *chymotrypsin-1*, two α-glucosidases, *peritrophin*, 2 chitinase proteins, *fibrillin-2*, *muscle protein 20*, and 7 uncharacterized genes. The OA-associated list also included *vegetative cell wallprotein-like (gpl-1)*, which exhibited higher expression in bees that flew further distances (mock, SBV, DWV and OA), than in DWV infected bees. Previous studies determined that *gpl-1* expression was greater in *Nosema ceranae* infected honey bees (78), and that octopamine signaling was also impacted (55, 56). While the bees in our study were not *Nosema* infected (S1 Fig), we used qPCR to confirm that bees expressed higher *gpl-1* levels when injected with OA or EP (LMM, *p* = 2×10^-7^ and 4×10^-8^, respectively); data from additional experiments (i.e., exp-1, n = 61 and exp-2, n = 69; Data S1).

We also examined expression of a subset of OA-associated genes including OA receptors (i.e., *oα-1R*, *oβ-1R*, *oβ-2R*, *oβ-3R, and oβ-4R*), enzymes involved in OA synthesis (*tdc* and *tβh*), and genes associated with Oβ-2R activation (*adenylyl cyclase* (*ac*), *protein kinase A* (*pka*), and CREB-associated protein encoding genes) (Data S9 and S5 Fig). No sample had detectable levels of *oβ-4R; oβ-1R* and *oβ-3R* were expressed at very low levels in most samples (Supplementary Table S9; Figure S5). Although *oα-1R* transcripts were detected in some samples, expression levels were consistently lower than those of *oβ-2R*. Expression of *tdc*, *tβh*, *oβ-1R*, *crebA*, and *creb-binding isoform X1* was generally higher in SBV infected honey bees, and greater *tdc* and *tβh* expression was confirmed via qPCR (exp 1-3, n = 336) (Fig 3, Data S1 and S4). Honey bees that were simultaneously DWV infected and treated with OA expressed higher levels of *oβ-2R*, *oα-1R*, and *creb binding isoform X2* than untreated DWV infected bees (S5 Fig); greater *oβ-2R* expression was also confirmed by qPCR (LMM, n = 336; *p* = 9×10^-4^; Data S1). Expression of *oa-1R* and *tyrR* was assessed via qPCR in only one experiment since expression was generally low regardless of treatment (exp-3, n = 69; Data S1). SBV infected bees had greater expression of *oa-1R* (*p* = 0.017), and SBV infected bees co-injected with EP had significantly lower expression of both *oa-1R* and *tyrR* (*p* = 0.0006 and 0.04, respectively).

## Discussion

The interactions between virus infections, host immune and stress responses, and organismal health are complex. Insects rely on OA as a neurohormone and neuromodulator to regulate physiological processes and behaviors (50, 52, 79–81), and our previous studies indicated that viruses differentially impacted honey bee octopamine signaling and flight performance. Therefore, we used honey bees as a model system to further investigate these interactions at organismal and transcriptional levels.

As previously described and shown herein, inapparent (or asymptomatic) DWV infections negatively impact honey bee flight performance (i.e., flight distance, duration, and peak flight speed). Evaluation of transcriptional responses to DWV infection suggest that DWV-associated flight impairment may be partially explained by lower expression of genes involved in energy production, which were elevated in OA treated honey bees (Fig 5, S5 Fig, Data S7-S8). These results are supported by previous studies that showed DWV infected honey bees expressed lower levels of G-coupled protein receptor encoding genes, as well as other genes involved in metabolism (82–84). Previously, we demonstrated that the flight performance of SBV infected honey bees was enhanced and that these bees expressed more *oβ-2R* (33). Herein, we showed that SBV-associated flight enhancement was reduced when OA signaling was inhibited by EP, an OA receptor antagonist (Fig 2 and 4). We expected that SBV infected bees would have compounding, enhanced flight performance when treated with OA, but unexpectedly, OA supplementation negatively impacted flight performance (Fig 2). To further investigate this physiological response, we examined the expression of key genes in the octopamine pathway and determined that in addition *oβ-2R*, SBV infected honey bees expressed greater levels of *tdc* and *tβh* (Data S1). Expression of *tdc* and *tβh* was lower in SBV infected bees that were treated with OA (Data S1 and S4, Fig 3-4). Reduced expression of these genes may limit OA synthesis, OA signaling, reduce metabolic activity, and result in less available energy for flight (50). Therefore, the reduced flight performance of SBV infected bees treated with OA may be explained by a feedback loop that regulates intracellular OA concentrations, though future studies that measure OA levels are needed to validate this hypothesis.

Depending on OA concentration and specific pathogen challenge, OA may enhance or inhibit immune function in many invertebrates, though mechanistic studies have been primarily restricted to two model organisms (57, 58, 85–87). In *Drosophila*, OA treatment impacts intracellular OA concentration, in a hypothesized feedback mechanism that regulates neurohormonal signaling (62, 88, 89). In *Caenorhabditis elegans,* OA pathways are hypothesized to be constitutively active, but downregulated upon pathogen infection to enable an effective immune response while limiting immune activity in the absence of pathogen challenge (85). Transcriptomic analysis of honey bees infected with a model virus (Sindbis) that were co-injected with double-stranded RNA, a virus-associated molecular pattern, had greater expression of an OA receptor (*oβ-2R*) (72). Results presented herein indicate that OA treatment of DWV infected honey bees alters expression of several genes associated with immune function including *apidermin*, *defensin*, and *melittin* (90–93). Additionally, DWV infected bees with OA treatment had reduced expression of an odorant binding protein that was previously associated with SBV infection, indicating that it may be involved in honey bee antiviral defense (94). Together, these results highlight virus-specific transcriptional responses in honey bees that may impact virus dynamics and flight performance during commonly occurring coinfections (33, 95, 96).

Multiple OA receptors have been identified in honey bee flight muscles and neural tissues including OαR1, Oβ-2R, Oβ-1R, and Oβ-3/4R (47). OA receptors have also been detected on invertebrate immune cells (57). In addition to their involvement in flight, OA receptors are also involved in honey bee hygienic, communicative dance, and thermoregulatory behaviors (47, 59, 63, 67, 97, 98). Typically, around 12 days post-emergence honey bees change from nurse bees, which care for developing brood, to pre-forager bees and undergo physiological changes required for flight (99–101). As honey bees age and become foragers, OA levels in the brain increase (51, 61, 102). Honey bees infected with the microsporidian parasite, *Nosema*, have greater OA levels than uninfected bees (55, 56) and are prone to precocious foraging (103), though *Nosema* levels did not appreciably impact honey bee flight capability (34). These findings are in-line with SBV-induced flight enhancement that correlated with higher *oβ-2R* expression levels (33). Previous laboratory-based studies demonstrated that precocious foraging may be induced by OA supplementation (51).

Honey bees are exposed to environmental OA sources including citrus blossom nectar, which may contain tyramine and OA at concentrations up to 256 µM (104–107). In addition, acaracides that are OA receptor agonists including amitraz and chlordimeform are commonly used in agriculture (108). Amitraz is commonly used to mitigate *Varroa* mite infestations in honey bee colonies (109). *Varroa* mites with amitraz-resistant Oβ-2R mutations have been reported (110, 111). Amitraz has generally been considered safe for honey bees since honey bee OA receptors are divergent from *Varroa* mite OA receptors (i.e., 48% identical at the amino acid level) (109, 112). However, honey bees exposed to amitraz had higher mortality when co-stressed via infection with a model virus (Flock House virus) (113). These results in conjunction with our findings, linking virus infection with the OA stress response pathway, may indicate previously unrecognized impacts at the organismal level. *Varroa* mites vector many honey bee viruses and infested colonies often have coinfections and greater average virus levels (114). Therefore, application of miticide to reduce *Varroa* and control vectored viruses, or the accumulation miticide metabolites in wax or other hive matrices, may inadvertently harm co-stressed, virus infected honey bees (113, 115, 116). Collectively, the data presented herein demonstrate that the OA pathway is a key factor in mediating virus-specific effects on honey bee flight performance, which may in turn influence viral transmission and colony health.

## Materials & Methods

Key Resources Table.

| Key Resources Table |  |  |  |
| --- | --- | --- | --- |
| Reagent type (species) or resource | Designation | Source or reference | Identifiers |
| epinastine-hydrochloride >98% | epinastine (EP) | Tokyo Chemical Industry | $C_{16}H_{15}N_3 \times HCl$ |
| octopamine-hydrochloride 99% | octopamine (OA) | ThermoScientific | $C_8H_{11}NO_2 \times HCl$ |
| gene (*Apis mellifera*) ribosomal protein L8 | *rpl8* | NCBI GeneID: 410188 | GB48810 |
| gene (*Apis mellifera*) octopamine β-2 receptor | *o\beta-2R* | NCBI GeneID: 412896 | Octbeta2R, GB49696, Octbeta2 |
| gene (*Apis mellifera*) tyrosine decarboxylase | *tdc* | NCBI GeneID: 410947 | Tdc1, GB55831 |
| gene (*Apis mellifera*) tyramine β-hydroxylase | *t\beta*h | NCBI GeneID: 751102 | AmTbH, GB16337, GB53665 |
| *Apis mellifera* transcriptome data | honey bee | NCBI SRA: PRJNA1414634 | Illumina sequencing of RNA obtained from virus-infected honey bees no- or post-flight, Jan 23 '26 |

### Honey bees

Honey bees (*Apis mellifera,* primarily *carnica*), were maintained with standard apicultural practices at Montana State University’s Horticulture Farm. Each experiment was performed with honey bees from different colonies to obtain robust results from genetically diverse bees (Data S1). Frames with newly emerging bees were collected one day prior to each experiment and maintained at 32^◦^C in a laboratory incubator overnight. Young, age-matched (∼24 h post-emergence), female adult bees were utilized for all experiments. Bees were cohoused in modified deli containers (n = 10 per house), maintained at 32°C in an incubator and fed 50% sucrose syrup and water *ad libitum* (72). All honey bees were fed only sucrose syrup, unless octopamine (OA) and/or epinastine (EP) oral treatment was otherwise specified. OA was administered at a concentration of 10 mM and EP at 4 mM in fresh sucrose syrup daily, and honey bees were given unrestricted access to the supplemented sucrose syrup for the duration of the experiment.

### Virus stock preparation

Deformed wing virus (DWV) and sacbrood virus (SBV) inoculums were propagated in white-eyed pupae as previously described (33). In brief, pupae were injected with 2 µL of Tris-HCl (pH 7.5) containing either 4.0×10^6^ DWV RNA copies or 3.5 x 10^4^ SBV RNA, between the 2^nd^-3^rd^ abdominal tergites using a Harbo syringe and a disposable borosilicate needle made from modified capillary tubes. Pupae were maintained in individual wells in a 24-well plate with a piece of Whatman filter paper at the bottom of each well to prevent pupal tissue damage. In each plate, four wells were filled with sterile water for humidity, at 32°C for 7 days. Pupae were then homogenized in a 2 mL microfuge tube with a sterile, metal, 3 mm bead in 1 mL 1X PBS (pH 7.4) using the TissueLyser II (Qiagen) at 30 Hz for 2 min. Homogenates were centrifuged at 14,000 x g for 15 min at 4°C and supernatant was transferred to a fresh tube. The virus-containing lysates were filtered through a 0.45 µm filter, then a 0.22 µm filter to remove large particles and non-viral microbial contaminants, respectively. RNA was isolated from the virus-containing filtrate (20 µL) using TRIzol (Thermo) according to manufacturer’s instructions. RNA was quantified using a NanoDrop 2000c Spectrophotometer, cDNA was synthesized with reverse transcriptase, and virus RNA copies (including both genomes and transcripts) were quantified from cDNA using qPCR. The inoculum was tested for copurifying/contaminating viruses via PCR (Data S2, S2 Fig). RNA copy concentration in the purified virus stock was quantified relative to a standard curve using qPCR (Data S3). DWV genome consensus sequences were assembled from short-read sequencing data obtained from sequencing RNA isolated from the inoculum (DWV-lab 2024; GenBank PV821422) and SBV sequencing data (sacbrood virus isolate MT 2023; GenBank PV788228.1). Virus inoculum aliquots were stored at –80°C until each experiment.

### Experimentally introduced honey bee virus infections

Honey bees were infected with DWV using previously described methods (72). In brief, age-matched bees were cold anesthetized at 4°C for 10 min and infected via intra-thoracic injection using a Harbo syringe with 3.5 x 10^4^ DWV RNA copies, 3.5 x 10^4^ SBV RNA copies, or coinfected with 3.5 x 10^4^ DWV RNA copies and 9.1×10^3^ SBV RNA copies. If octopamine (OA) and/or epinastine (EP) treatment was administered via injection, the injection inoculum also contained 10 mM OA and/or 4 mM EP so that each bee only experienced one injection to reduce injection-associated stress. Each inoculum dose was suspended in 2 µL10 mM Tris HCl buffer, pH 7.5. Buffer injected/ mock infected bees were injected with 2 μL buffer. The DWV doses utilized in this study were similar to previous studies and resulted in virus levels commonly found in naturally infected bees (8, 34, 37, 38, 117, 118). As previously described (33), bees had #000 steel washers affixed to their thoraces using rubber cement immediately after injections; washers had an outside diameter of 0.19 cm and weighed 4.3 mg, less than half of an average pollen load (3).

### Flight mill

The flight mills used in this study were based on flight mills used in previous honey bee studies (33, 46, 119, 120) and included a magnet to tether bees from the affixed steel washer on each thorax, which enabled their removal from flight mills with minimal damage. Flight assays were performed as previously described (33). In brief, bees 72 hpi (i.e., 4-days post eclosion) were immobilized via incubation at 4°C for ∼12 minutes and tethered to flight mills; counterweights were adjusted for each bee. Temperature and humidity were maintained the same for each experiment (23°C and 31±4% relative humidity). Flight was instigated by tapping the flight arm downward and was re-initiated up to three times before bees were considered ‘exhausted’ and the data collected was representative of ‘total flight’ capability (Data S1). Bees were stored at – 80°C immediately after flight. Flight distance, duration, and speed calculations were based on redlight detector counts which were recorded on an attached Raspberry Pi as previously described (33). Active flight duration was calculated as the sum time moving >5.5 cm (one encoder count) (Data S1). Flight stop count was defined as any stop that exceeded 1.5 seconds. Honey bees may stop to seemingly engage in grooming behaviors, stops were counted if the bee re-initiated flight (33). DWV infected, EP injected, SBV infected bees fed OA or EP or injected with OA, or SBV infected bees fed both OA and EP had fewer stops than mock infected bees (Data S4).

### Honey bee RNA isolation

RNA was isolated from the abdomen of each individual honey bee since abdomens were representative of the entire bee (33). Each abdomen was individually homogenized in sterile water (300 μL) with one sterile steel bead (4.5 mm) using a Qiagen TissueLyser for 2 min at 30 Hz. Lysates were centrifuged at 4°C for 5 minutes at 7,500xg. RNA was isolated from the supernatants using equal volumes of TRIzol reagent according to manufacturer’s instructions. The concentration and quality of RNA samples were assessed on a ThermoFisher Nanodrop 2000 spectrophotometer. RNA was stored at −80°C.

### Reverse transcription / cDNA synthesis

Reverse transcription reactions were performed by incubating 2 μg RNA, 200 U M-MLV reverse-transcriptase and 500 ng random hexamer primers in 25 μL reactions for 1h at 37°C, according to the manufacturer’s instructions. cDNA was diluted with sterile water (1:2) and 2 μL were used for polymerase chain reactions (PCR).

### Polymerase Chain Reaction

Since bees were obtained from colonies that are subject to natural infections (37, 72), PCR was performed to test for preexisting virus infections. Specifically, 2 μL cDNA template was combined with 10 pmol of each forward and reverse primers (Data S2) and amplified with ChoiceTaq polymerase according to the manufacturer’s instructions using the following conditions: 95°C for 5min; 95°C for 30s, 57°C for 30s, 72°C for 30s, 35 cycles, followed by final elongation at 72°C for 4min. PCR products were assessed by gel electrophoresis (1.5% agarose with SYBR safe) and visualized using a Syngene U:Genius 3 imaging system (S1 Fig). PCR products were previously verified by sequencing (37). Analysis of pooled cDNA samples from mock infected honey bees from each of these experiments revealed that many bees had preexisting DWV and SBV infections, therefore qPCR was used to assess their abundance in all individual samples (Data S1). The qPCR primers used in this study to target DWV, bind to both DWV-A (GenBank AY292384) and DWV-B (GenBank MN565036), in addition to numerous other DWV sequences on NCBI including the consensus sequences of the DWV inoculum as part of this study (Datal S3).

### Quantitative PCR

Quantitative PCR (qPCR) was used to assess virus abundance and relative abundance of honey bee transcripts. All qPCR reactions were performed in triplicate with 2 μL of 1:2 diluted cDNA template. Each reaction contained 1× ChoiceTaq Mastermix, 0.4 μM each forward and reverse primer, 1× SYBR Green (Life Technologies), and 3 mM MgCl_2_ for a total volume of 20 μL per reaction. A CFX Connect Real Time instrument (BioRad) was used for the following: pre-incubation 95°C for 1 minute followed by 40 cycles of 95°C for 10s, 58°C for 20s, and 72°C for 15s, with a final melt curve analysis at 65°C for 5s to 95°C. To quantify virus copies in each sample, plasmid standards (i.e., virus-specific PCR amplicons cloned into the pGEM-T (Promega) vector and sequence verified) for each virus were used as templates, ranging from 10^3^-10^9^ copies per reaction for each standard curve as previously described (Data S3) (33). The honey bee gene *rpl8* was used as a housekeeping gene for each sample for comparison and for relative fold change calculations (Data S1-S2). Sterile water containing no DNA template was used for negative controls. qPCR specificity was verified by melt point analysis, gel electrophoresis, and sequencing (37). The starting quantity (SQ) of cDNA template was calculated for each honey bee sample as previously described (33). For each honey bee sample, the starting quantity (SQ) of cDNA template (representing 80 ng total RNA) was calculated based on the standard curve and subtracting the average SQ of the no template control reaction (<600 copies). Samples with virus RNA copies below the limit of detection for qPCR (<1000 copies / 2 µg RNA) were listed as 0 (Data S1). Virus abundance was reported as virus RNA copies, including genomes and transcripts, per 2 µg total RNA (i.e., per RT reaction) and ranged up to 7×10^10^ virus RNA copies / 2µg RNA. Whole bee data was estimated based on the previous estimate of 138 µg RNA per whole bee (33). Comparison of virus abundance levels using estimated copy numbers enabled inclusion of values of 0. Using the ΔΔCt method to calculate virus levels in bees with no detectable virus would inaccurately result in ‘missing’ data and consequently skew model interpretations. Relative host gene expression was calculated using the average ΔΔCt method where ΔCt was determined by subtracting *rpl8* Ct from the gene of interest Ct. Average ΔCt was calculated by the mean ΔCt of the buffer-injected bees in each experiment, which harbored the lowest virus levels. The ΔΔCt was calculated by subtracting the average control ΔCt and fold change was determined by the equation 2^-ΔΔCt^ (Data S3).

### RNA sequencing and analysis

Honey bee samples with low/no preexisting infections (n = 3-5) were selected per treatment from previously published data (33) (n=36 total) (Data S5). RNA from each sample was DNase treated using Qiagen RNeasy columns. RNA samples were sent to the Roy J. Carver Biotechnology Center at University of Illinois; quality was assessed with an Agilent 3500 fragment analyzer. Libraries were prepared via Watchmaker polyA RNA-Seq prep kit according to manufacturer instructions. Libraries were sequenced on an Illumina NovaSeq X Plus 25B (paired 150 nt reads), approximately 3.2 billion total paired end reads were obtained. Sequencing quality of reads representing individual samples using FastQC, and filtering of the comprehensive dataset was done using MultiQC versions 0.12.1 (121). Illumina adaptors were trimmed and low quality reads were omitted when Q<30 with BBduk from BBTools version 39.26 (122). Reads were normalized with shrinkage estimations for dispersion using DESeq2 l (correction with q-value<0.05) (123). Sequencing libraries were aligned to the *Apis mellifera* genome, Amel_HAv3.1 (i.e, GCA_003254395.2_Amel_HAv3.1_genomic.fna, downloaded from NCBI 2019–02-07) (73, 74). Sequencing data from individual samples were uploaded to the NCBI short read archive (SRA) (BioProject ID: PRJNA1414634). Genes were considered differentially expressed if log_2_ fold change of transcript abundance had adjusted p-values <0.05. Venn diagrams were produced using the eulerr package in R (124).

For virus quantification, trimmed paired end read files were aligned to a nonredundant version of the Holobee database (i.e., https://doi.org/10.15482/USDA.ADC/1255217) (125) which considered 32 potential virus alignments, using HISAT2 version 2.2.1 (126). Default parameters were used during alignment to allow multimapping of reads. Samples with unexpected infections were omitted from analyses (n = 6/36; Data S5). Alignment files were converted to compressed alignment files and sorted using samtools version 1.21 and unmapped reads were discarded (127). The samtools built-in idxstats command was used to quantify virus abundance in sorted alignment files and generate read counts. Read counts of each trimmed FastQ file were quantified using the Seqkit stats command (128). Computational efforts were performed on the Tempest High Performance Computing System, operated and supported by University Information Technology Research Cyberinfrastructure (RRID:SCR_026229) at Montana State University.

### Statistics

Statistical analyses were performed in R 4.3.3. Log-transformations and non-parametric tests were performed as necessary. Specifically, all relative gene expression data was log_2_ transformed and all virus abundance data were log_10_(1+) transformed, which enabled inclusion of 0s. All mixed models were assessed using the lmer4 package (129). Assumptions for linear models were verified by diagnostic plots and histograms using base R functions and the performance package (130, 131). All models included experiment ID as a random effect and where relevant, virus injections and flight mill IDs were also included as random effects (Data S4). Models were compared using corrected Aikike information criterion (AICc) and Bayesian information criterion (BIC). Models were assessed via maximum likelihood (ML) when random effects varied and via restricted maximum likelihood (REML) when fixed effects varied. We reported only results from the best model for each analysis, but other considered models were included in supplementary material (Data S4). For each linear mixed model with reported estimates, *p*-values were estimated from *t*-statistics via Satterthwaite’s method (132).

### Figures

All figures were generated through GraphPad Prism software for windows using version 10.6.0, in RStudio using version R 4.3.3, in BioRender, and Adobe Illustrator.

## Acknowledgments

The authors would like to thank Dr. Mark Jankauski and Larry Jankauski for help with design and construction of the flight mill data recording system, Darion Christiansen for scripting, Boone Jones for assistance with analysis of sequence data, and members of Montana State University for valuable feedback on this manuscript.

## Funding

This work was supported by National Science Foundation (NSF) Integrative Organismal Systems (IOS), Physiological and Structural Systems (PSS) Cluster, Symbiosis, Infection and Immunity (SII) Program Award (2348112). In addition, Naomi Kaku was partially supported by a National Center For Advancing Translational Sciences of the National Institutes of Health Award (TL1TR000422). The funders had no role in study design, data collection and analysis, decision to publish, or preparation of the manuscript.

## Author Contributions

N.G.K. and M.L.F. designed research; performed research; analyzed data; and wrote the paper. M.L.F. acquired funding, provided resources, supervised the research, and administered the project.

## Competing Interest Statement

N.G.K and M.L.F. declare that they have no competing interests.

## Data and materials availability

All data are available in the main text or the supplementary materials. Additional information and requests should be directed to the corresponding author, Michelle Flenniken.

## Notes

### Competing Interest Statement

The authors have declared no competing interest.

### Summary of Updates

This version of the manuscript has been revised to update the following: minor editorial changes and corrections.

