## Supplemental Figures for "Virus specific impacts on honey bee flight performance are mediated by the octopamine pathway"

### Figure S1. Pathogen testing of virus inoculum and honey bee samples

(A) Filtered virus inoculums DWV, SBV, and DWV+SBV (labeled A, B, or C, respectively) were tested for other common viruses including acute bee paralysis virus (ABPV), Apis mellifera filamentous virus (AmFV), Andrena bee-associated virus-1 (AnBV), black queen cell virus (BQCV), chronic bee paralysis virus (CBPV), deformed wing virus (DWV), Israeli acute paralysis virus (IAPV), Kashmir bee virus (KBV), Lake Sinai viruses 1-4 (LSV), and sacbrood virus (SBV) using virus-specific polymerase chain reaction (PCR) and analyzed by gel electrophoresis; positive (+) and negative (-) controls. (B) Pathogen diagnostic PCR was performed using pooled mock-infected honey bee cDNA from experiments 1-3 (labeled 1-3) as template determined that bees had preexisting DWV (experiments 1-3) and SBV (experiments 1-3). All samples were tested for DWV and SBV via qPCR (Table S1). All experiments were negative for all non-viral pathogens including *Ascosphaera apis* (*Aa.*), *Lotmaria passim* (*Lp.*), *Melisococcus plutonius* (*Mp.*), *Nosema ceranae* (*Nc.*), and *Paenibacillus larvae* (*Pl.*); cDNA quality was assessed via amplification of the honey bee housekeeping gene, *rpl8*. Although no positive control was available for *A. apis* or *P. larvae*, the primers were utilized successfully in previous studies.

To illustrate this result in a single figure, cDNA from mock-infected individual bee samples from all experiments were pooled by experiment (i.e. n=12 from experiment 1, n=12 from experiment 2, n=12 from experiment 3) and PCR was repeated. The products of pathogen-specific PCRs using pooled cDNA (S), positive (+), and negative control (-) templates were analyzed by agarose gel electrophoresis

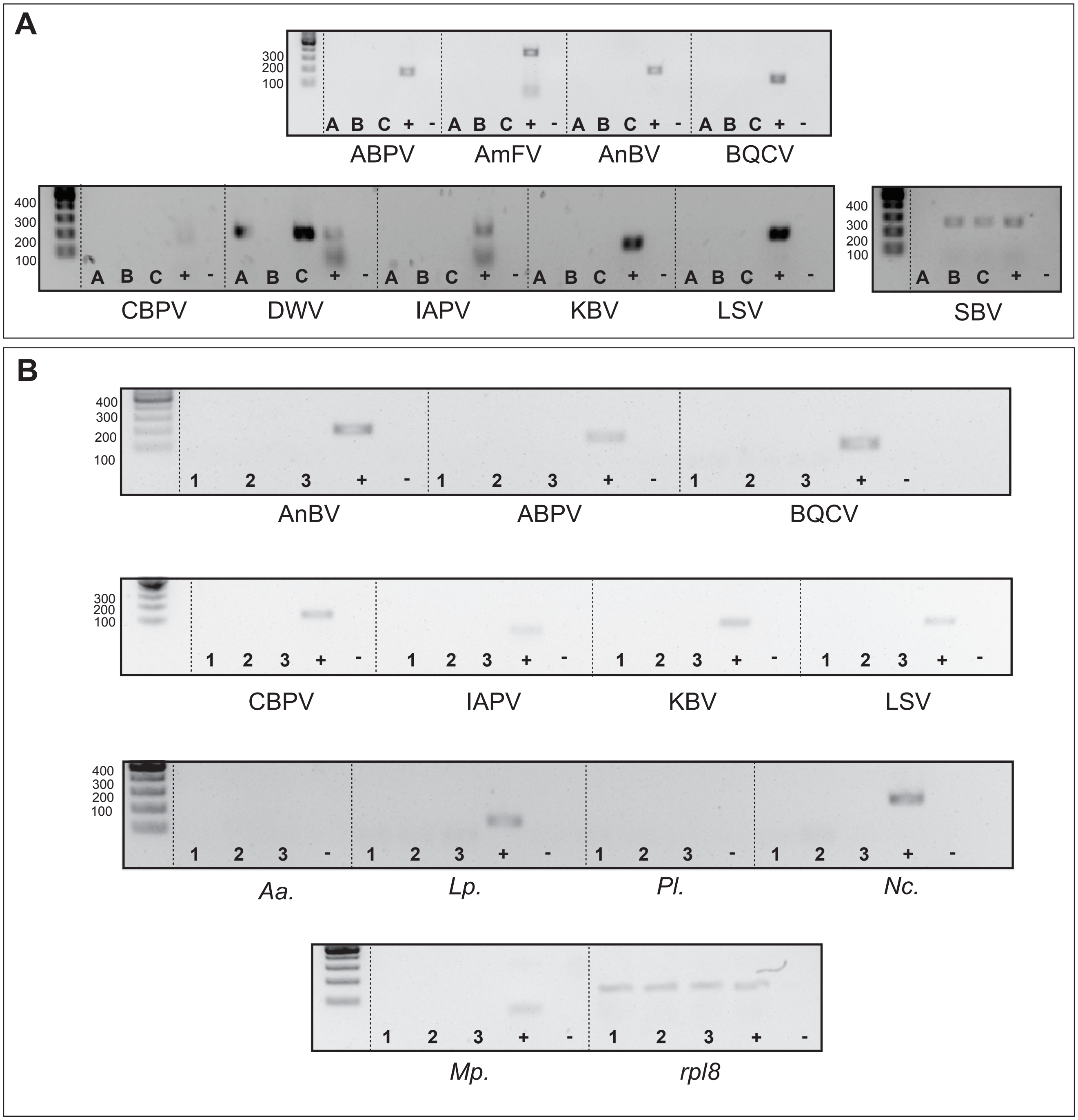

### Figure S2. Estimated flight duration and speed by virus and OA and/or EP treatment

To visualize the effects of virus infection and octopamine (OA) and/or epinastine (EP) treatment on honey bee (A-D) flight duration (minutes) and (E-H) peak flight speed (km/h), we compared estimated means using experimental data analyzed with linear mixed models (S1 Appendix Tables  Table S4). Each individual point represents the estimated marginalized means with bars indicating one standard deviation of the mean. Green points represent predictions when bees harbored high SBV levels (i.e., $10^8$ SBV RNA copies / 2 µg RNA) and orange points represent predictions when bees harbored high DWV levels (i.e., $10^8$ DWV RNA copies / 2 µg RNA). Blue points represent estimates for virus free bees. (A) SBV infected bees flew similar durations to uninfected bees, and all treatments resulted in similar or shorter flight durations than those fed sucrose only except OA fed bees, which flew for greater durations. (B) DWV infected bees flew for shorter durations than uninfected bees, but EP and OA treatment resulted in greater flight durations. (C-D) Observed flight durations where each point represents data collected from an individual bee (total n = 336) and the color scale represents virus levels. (E) SBV infected bees flew similar peak speeds than uninfected bees, but any OA or EP treatment resulted in lower peak speeds except for OA+EP injected and OA injected bees, which flew similar distances to those fed only sucrose. (F) DWV infected bees flew at slower speeds than uninfected bees. When DWV infected bees were fed or injected with OA, peak speeds were slightly higher. (G-H) Observed peak flight speeds where each point represents data collected from an individual bee with (G) SBV infections or (H) DWV infections.

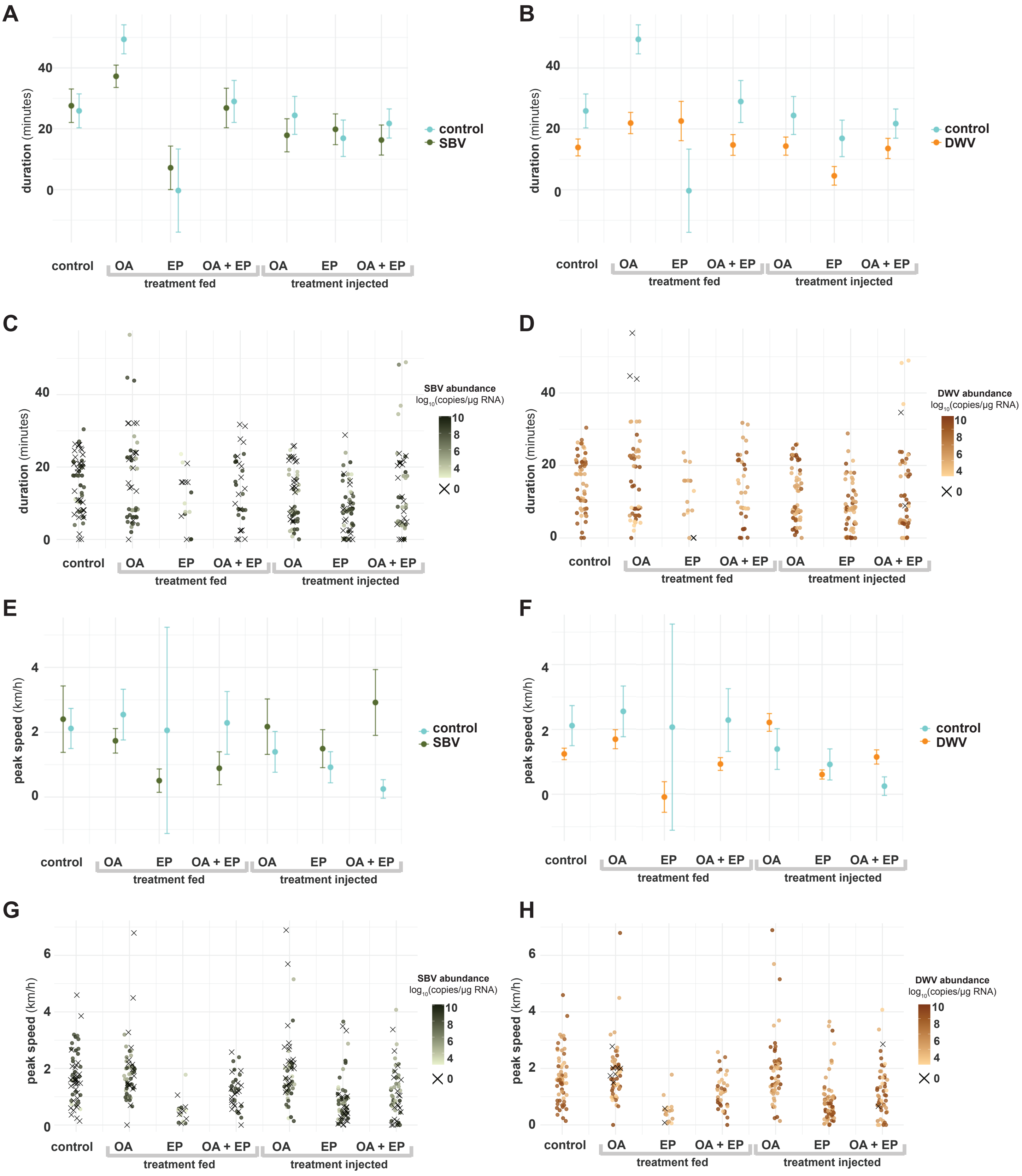

**Figure S3. DWV infected bees had a no relationship with *tdc* and treatment-specific relationships with *tβh***

To identify relationships between *tyrosine decarboxylase* (*tdc*) and *tyramine β-hydroxylase* (*tβh*) expression across virus infection and treatments, we evaluated relationships using a linear mixed effect model (SI Appendix Table S4). To visualize the data, we plotted all data points against treatment groups with either (A) *tdc* expression or (B) *tβh* expression. The background violin plots represent the 95% confidence interval, the individual data points represent individual bee data, the horizontal line in each violin represents the median for the treatment group. Ranges of DWV infection were included as a colorscale (i.e., darker orange representing higher DWV infection levels). If there is no effect of DWV infection, the darkest points would be evenly distributed above and below the control '0' fold change line. SBV levels were more strongly associated with *tdc* expression (Fig. 3).
(A) There was no difference in *tdc* expression by DWV abundance ($p$ = 0.83). (B) The expression of *tβh* was greater in DWV infected bees relative to uninfected bees ($p$<0.0001, SI Appendix Table S4). In addition, DWV infected bees fed OA, injected with EP, or injected with OA and EP exhibited reduced *tβh* expression ($p$-values <0.001, SI Appendix Table S4).

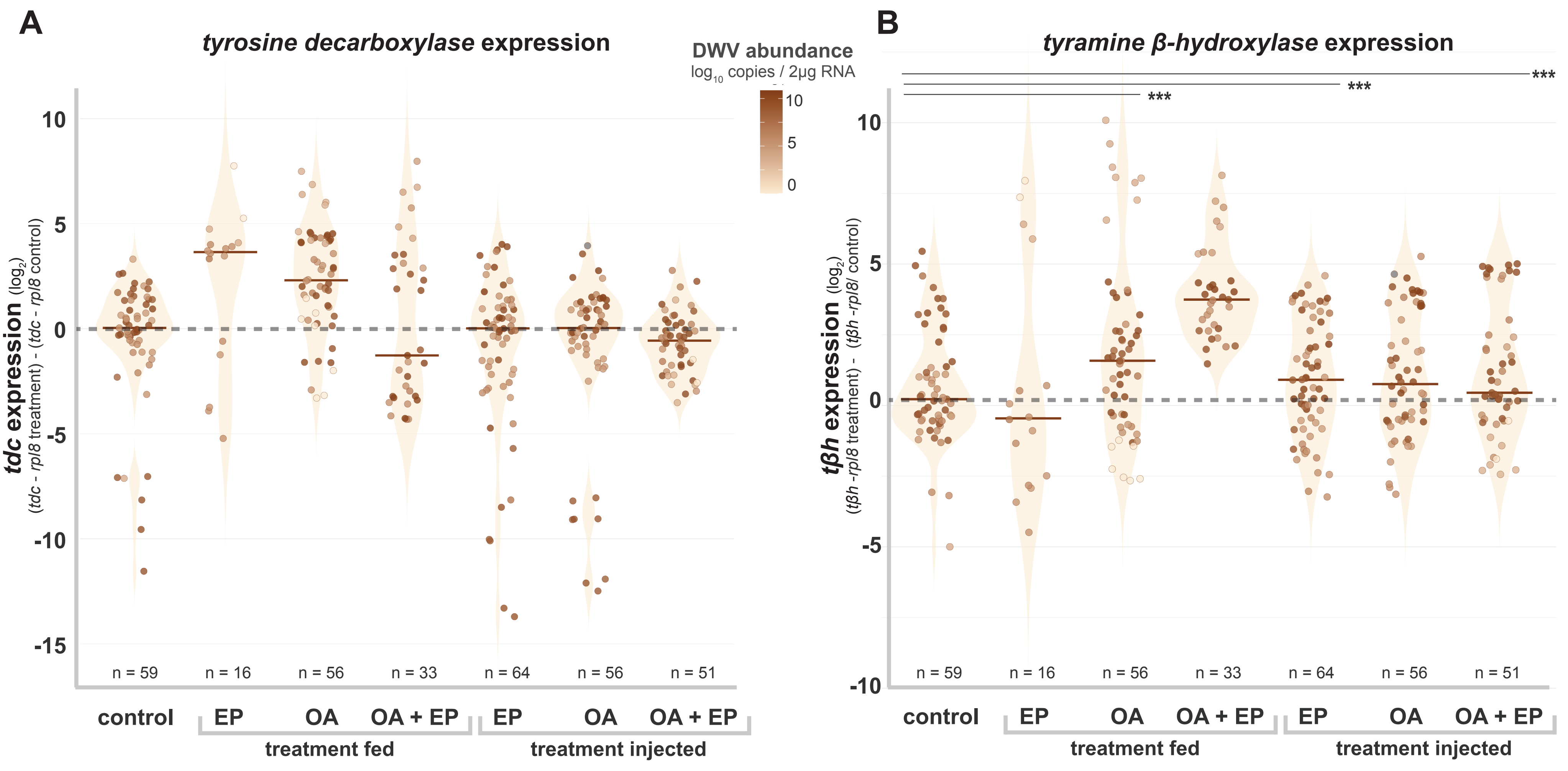

### **Figure S4. Expression of *tdc*, *tβh*, and *oβ-2R* are positively correlated**

The relationships between the expression of *tdc*, *tβh*, and *oβ-2R* in honey bees were evaluated using a linear mixed effect model that included *tdc*, *tβh*, and *oβ-2R* expression, and SBV abundance as fixed effects and experiment, injection, and treatment (i.e., OA and/or EP) as random effects. This model, which explained 95% of the data, determined that *tdc*, *tβh*, and SBV abundance were associated with increased *oβ-2R* expression (*p*-values <0.05, SI Appendix Table S4). Each data point represents data from an individual honey bee across three experiments (total n = 336; SI Appendix Tables S1 and S9).

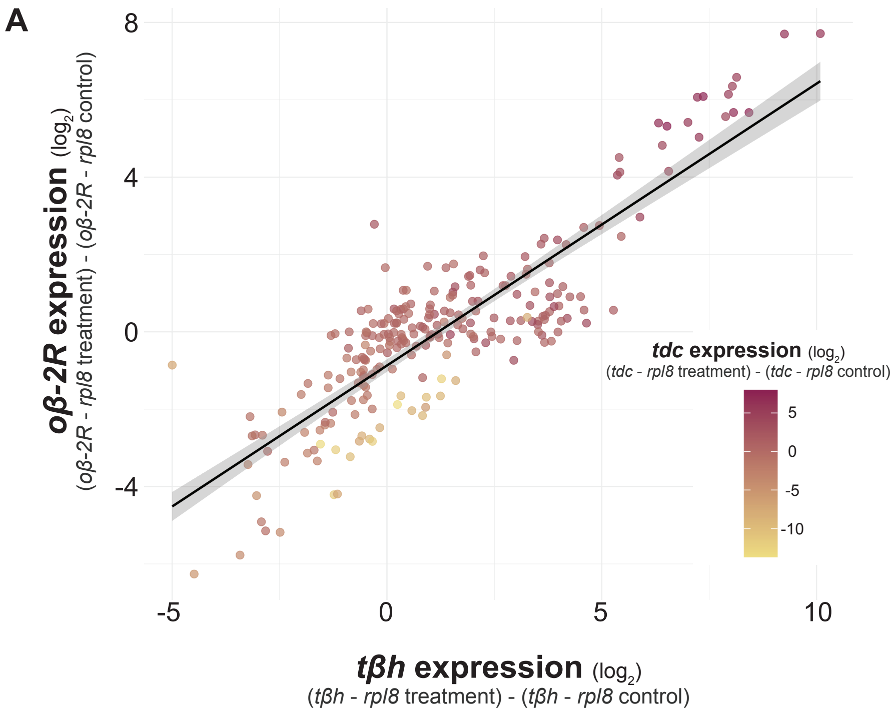

### Figure S5. Comparison of honey bee OA-associated gene expression

To identify potential differences in expression between different treatments and octopamine (OA)-associated genes, expression (as variance stabilized transformed values) between different treatments were evaluated. Specifically, the expression of creb subunits (A and B), protein Kinase A (PKA) subunits, OA receptors (*oβ-2R, oβ-3R, oβ-1R,* and *oα-1R*), *adenylyl cyclase* (*ac*) and the two precursor enzymes that convert tyrosine to tyramine and tyramine to OA (i.e., *tdc* and *tβh*, respectively). Each point represents data from an individual bee. Blue boxplots were mock infected with buffer injections, green boxplots were experimentally-infected with sacbrood virus (SBV), orange boxplots were experimentally infected with deformed wing virus (DWV), and pink boxplots were coinjected with DWV and OA (DWV+OA). Lighter colors indicate treatments that did not fly and darker color boxplots indicate treatments that flew (see Fig. 5). We compared the average expression of OA associated genes across treatment groups and determined that SBV infected bees that did not fly had the greatest differences in OA-associated gene expression relative to all other treatment groups (Complete data available in SI Appendix Table S9).

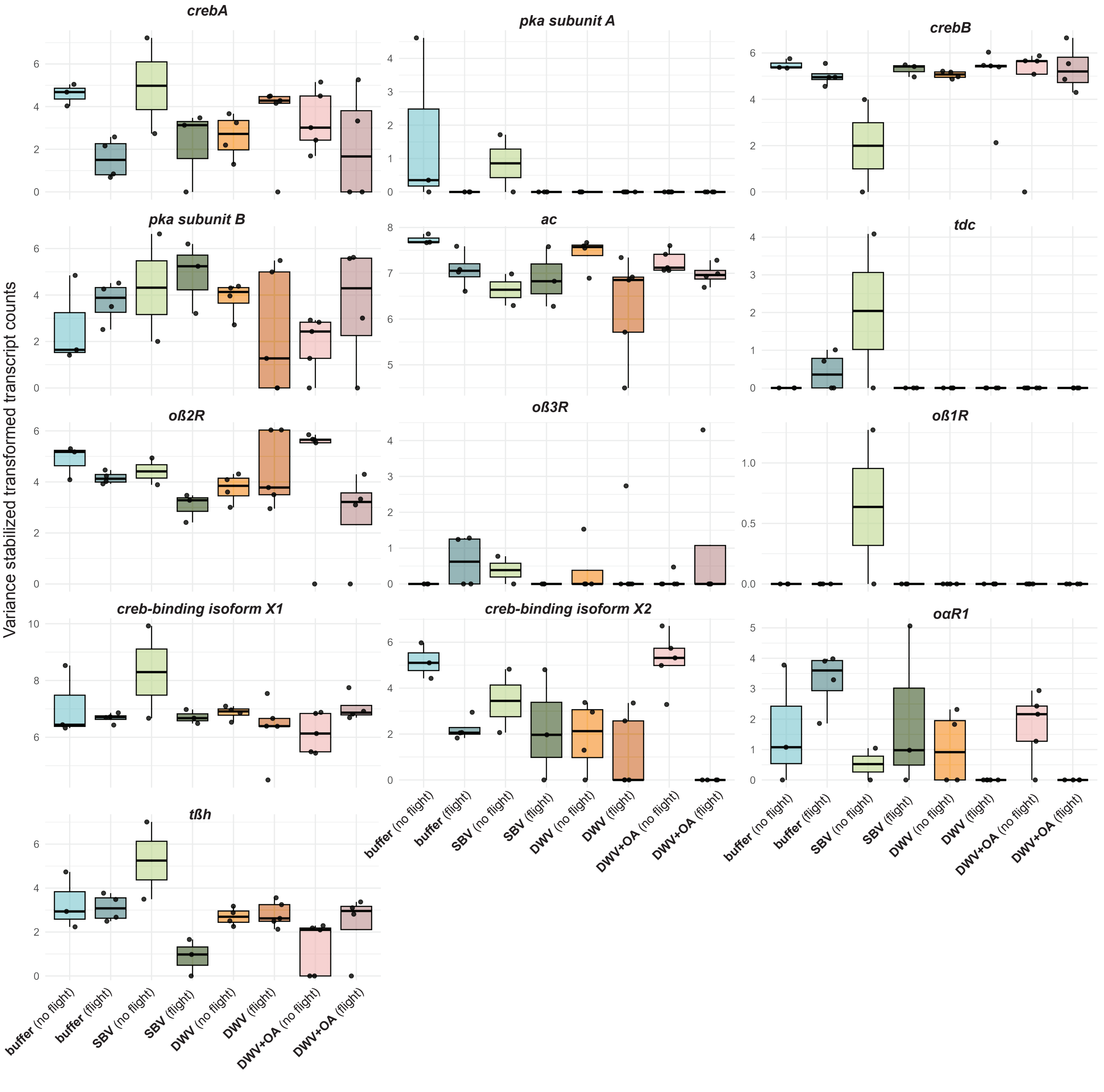
